# Novel inhibitors of TTLL12’s oncogenic potential that overcome suppression of ligation of nitrotyrosine to the C-terminus of detyrosinated α-tubulin

**DOI:** 10.1101/2023.12.28.573570

**Authors:** Amit Deshpande, Jan Brants, Christine Wasylyk, Onno van Hooij, Gerald W Verhaegh, Peter Maas, Jack A Schalken, Bohdan Wasylyk

## Abstract

Tubulin tyrosine ligase 12 (TTLL) is a promising target for therapeutic intervention since it has been implicated in tumour progression, the innate immune response to viral infection, ciliogenesis and abnormal cell division. It is the most mysterious of a fourteen-member TTL/TTLL family, since, although it is the topmost conserved in evolution, it does not have predicted enzymatic activities. TTLL12 seems to act as a pseudo-enzyme that modulates various processes indirectly. Given the need to target its functions, we initially set out to identify a property of TTLL12 that could be used to develop a reliable high-throughput screening assay. We discovered that TTLL12 suppresses the cell toxicity of nitrotyrosine (3-nitrotyrosine) and its ligation to the C-terminus of detyrosinated α-tubulin (abbreviated to ligated-nitrotyrosine). Nitrotyrosine is produced by oxidative stress and is associated with cancer progression. Ligation of nitrotyrosine has been postulated to be a check-point induced by excessive cell stress. We found that the cytotoxicities of nitrotyrosine and tubulin poisons are independent of one another, suggesting that drugs that increase nitrotyrosination could be complementary to current tubulin-directed therapeutics. TTLL12 suppression of nitrotyrosination of α-tubulin was used to develop a robust cell-based ELISA assay that detects increased nitrotyrosination in cells that overexpress TTLL12 We adapted it to a high throughput format and used it to screen a 10,000 molecule World Biological Diversity SET^TM^ collection of low-molecular weight molecules. Two molecules were identified that robustly activate nitrotyrosine ligation at 1 μM concentration. This is the pioneer screen for molecules that modulate nitrotyrosination of α-tubulin. The molecules from the screen will be useful for the study of TTLL12, as well as leads for the development of drugs to treat cancer and other pathologies that involve nitrotyrosination.

## Introduction

TTLL12 is an unconventional member of a family of factors that modify α-tubulin post-translationally. The family is composed of tubulin tyrosine ligase (TTL) and 13 related tubulin tyrosine ligase like (TTL) factors [recent reviews: (1, 2)]. TTL binds to the C-terminus of α-tubulin (3) and religates tyrosine that has been removed by the detyrosinases, SVBP/VASH (4, 5) and MATCAP (6). Nine TTLLs catalyse tubulin glutamylation (7, 8) and four glycylation (9) (8). The remaining TTLL, TTLL12, appears to be a pseudo-enzyme (10). TTLL12 has SET-like and TTL-like domains but no apparent consequential activity, as either a methylase, tyrosine ligase, glutamylase or glycylase (8-10). However, TTLL12 has indirect effects on tubulin detyrosination, acetylation and methylation, as well as histone methylation (10, 11) [preprint: (12)]. This poorly understood TTLL merits further study, and the development of tools to probes its functions, especially given its wide implication in pathologies and cellular functions.

We originally selected TTLL12 as a potential target for drug development from our multi-omic dissection of human tumours and bioinformatic predictions of enzymatic activity. We found that TTLL12 is expressed in the proliferating layer of benign prostate and its expression increases during cancer progression (11). This overexpression in cancer could generate serum autoantibodies that recognise TTLL12 specifically in prostate cancer patients (13). TTLL12 expression and alterations have been linked to prognosis of various cancers, including prostate (14), stomach (15), lung adenocarcinoma (16), ulcerative colitis (17), ovarian cancer (18), and keloids in Chinese patients (19). A novel transcript isoform has been identified in human cancers (20). TTLL12 may contribute to tumour progression through effects on chromosomal ploidy (11), mitotic duration (10) and microtubule dynamics [preprint: (12)]. Interestingly, TTLL12 is very highly conserved in evolution (10, 21) and the only related protein in rice affects the dynamics and orientation of microtubules (22).

TTLL12 is also implicated in the innate immune response to viral infection. It acts as a negative regulator of RNA-virus induced type-I IFN expression through protein-protein interactions, rather than protein modifications (23). Repression of the innate immune response could explain different observations. TTLL12 is differentially expressed between HIV positive patients that have or have not been treated with highly active antiretroviral therapy, as well as healthy controls (24). TTLL12 activation improves lentiviral vector yields in HEK293 cells, which was found using genome-wide activator- and inhibitor-CRISP library screens to identify perturbations that improve viral yield (25). Downregulation of TTLL12 gene transcription by the vitamin-D receptor / vitamin D has been proposed to alleviate the severity of SARS-CoV-2 infection (26). TTLL12 genetic variants are linked to the heritable gut microbiome induced by Salmonella Pullorum in chicken (27). However, the immune response is not obviously linked to other observations. TTLL12 is: a target for serum autoantibodies after acute angle closure glaucoma attack (28), a candidate gene for triglycerides in mice (29), and a gene that is differently expressed during lactogenic differentiation of buffalo mammary epithelial cells (30). Interestingly, TTLL12 is required for the formation of primary ciliary axonemes, which are critical sensory organelles in polarized epithelial cells [preprint: (12)].

Amongst TTLL12’s functions that can be targeted for drug development, we chose inhibition of tubulin retyrosination for various reasons. Such anti-TTLL12 drugs could be complementary to current therapeutic agents that target microtubules [review: (31)]. They could target pathologies that involve dysregulation of α-tubulin retyrosination, such as cancer, neurodegenerative diseases, cardiovascular diseases and inflammatory disorder [reviews: (2, 32) (33)]. They may also target proteins other than α-tubulin that are tyrosinated by TTL (13), as well as stress pathways that are proposed to involve ligation of nitrotyrosine (3- nitrotyrosine) to detyrosinated α-tubulin (34). Nitrotyrosine ligation may lead to microtubule dysfunction, as reported by some (34, 35) but not all (36) studies. Nitrotyrosine is formed by a reaction between tyrosine in proteins and nitric oxide-derived reactive nitrogen species, such as peroxynitrite. Nitrotyrosine formation is a marker of oxidative and nitrosative stress, and has been implicated in various pathological conditions, including inflammation, neurodegenerative disease, cardiovascular diseases, and cancer [reviews: (37) (38)]. Nitrotyrosine levels increase with progression of various cancers, including prostate (39, 40) and HNSCC (41). Nitrotyrosine, derived from degradation of nitrotyrosine containing proteins and ligation to α-tubulin, could be a “last” checkpoint leading to apoptosis, and escape from the checkpoint could lead to cancer (42, 43).

We report that TTLL12 inhibits both nitrotyrosine incorporation into α-tubulin and nitrotyrosine induced cell toxicity. Using a validated high throughput screen (HTS), we identified hit compounds that relieve nitrotyrosination suppression by TTLL12. This is an important step towards developing molecules that could be used to probe the functions of TTLL12 and ultimately treat cancer and other pathologies by targeting novel pathways.

## Materials and Methods

### Cell culture

**HEp-2.** Human Caucasian larynx carcinoma cells (HEp-2) were obtained from American type cell culture (ATCC) (Cat. No. CCL-23), cultivated in modified Eagle’s medium (MEM) supplemented with 10% fetal calf serum (FCS), 0.1mM non-essential amino acid, 1 mM Sodium pyruvate, 40 µg/ml gentamycin. **A549.** Human Caucasian lung carcinoma cells (A549) were obtained from ATCC (Cat. No. CCL-185), cultivated in Dulbecco’s Modified Eagle’s Medium (DMEM) Ham’s nutrient mixture F12 (1:1) supplemented with glutamax, 10% fetal calf serum (FCS), 0.1mM non-essential amino acid, 1 mM Sodium pyruvate, 40 µg/ml gentamycin. **PC3.** Human male Caucasian grade IV prostatic adenocarcinoma (PC3) derived from bone metastasis were obtained from ATCC (Cat. No. CRL-1435), cultivated in DMEM: HAM’s F12K (1:1), 10% FCS, gentamycin 40 µg/ml. **DU145.** Human male Caucasian prostate carcinoma cells (DU145) derived from brain metastasis were obtained from ATCC (Cat. No. HTB-81), cultivated in MEM (EAGLE), 10% FCS, Sodium pyruvate 1mM, 0.1mM non-essential amino acid, 40 µg/ml gentamycin. **RWPE-1.** Human male Caucasian prostate epithelial cells (RWPE-1) transformed with human papilloma virus, were obtained from ATCC (Cat. No.CRL-11609), cultivated in keratinocyte-serum free medium (Life Tech 17005-042), human recombinant EGF 5ng/ml, bovine pituitary extract 0.05mg/ml. **PWR-1E**. Human male Caucasian prostate epithelial cells (PWR-1E) transformed with Adenovirus 12 and SV40 DNA virus were obtained from ATCC (Cat. No. CRL-11611), cultivated in keratinocyte-serum free medium (Life Tech 17005-042), human recombinant EGF 5ng/ml, bovine pituitary extract 0.05mg/ml. **TTLL12 and TTL stable clones.** The clones were generated in the laboratory (10). Briefly, for TTLL12 stable clones TTLL12 cDNA from 22Rv1 prostate cancer cells, and for TTL stable clones, TTL from human fetal brain EST (ATCC Cat. No. MGC-46235) cloned into pSG5-Puro Fnt expression plasmid were transfected in HEp-2 cells using the BBS calcium phosphate method (Chen et al., 1997). Empty pSG5-Puro Fnt expression plasmid was used for control clones. Two days post-transfection, cells were passaged into selection medium containing 3 μg/ml puromycin, which was renewed every three days. 10-14 days post transfection, five positive clones per stable transfection were characterized by immunofluorescence and western blotting (WB), giving rise to TTLL12 clones A-E, TTL clones A-E and Control clones A-E.

### siRNA TTLL12

siRNAs used in this study include: non-coding control siRNAs: siControl (CONTROL® Non-Targeting siRNA #1, Dharmacon); siLuciferase (GL2 luciferase siRNA, Dharmacon); siSilencer (Silencer® Negative Control #1 siRNA, Ambion); siGFP (GFP-22 siRNA, Qiagen), TTLL12-specific siRNAs: siTTLL12_1: 5’-GAGUUCAUCCCCGAGUUUG-3’; siTTLL12_2: 5’-GGAACGAGCUGUGCUACAA-3’; siTTLL12_3: 5’- AAGGCCAUCUUCUCUUAAA-3’; siTTLL12_4: 5’-ACGCCGACAUCCUCUUCAA-3’; siTTLL12_5: 5’-GUAGCGGUGUCUCCUCUUU-3’ (Dharmacon); siTTLL12_6: 5’- GGUUGUUCGUGUAUGAUGU-3’ (Sigma-Proligo).

### Transient transfections

siRNAs were transfected with Lipofectamine (Invitrogen Cat No. P/N50470) according to the manufacturer’s protocol. In brief, 100,000 cells for HEp-2 and 50,000 cells for DU145 were seeded per well in a 12 well plate. After 18 h, cells were incubated for 3 h in OPTIMEM prior to transfection. Unless otherwise indicated, 10 nM siRNA was transfected per well (5 µl Lipofectamine/12 pM siRNA in OPTIMEM). Six hours after transfection, medium was changed to normal growth medium and incubated for the desired time period. Plasmids were transfected using Lipofectamine in 12 well plates as for siRNA (5 µl lipofectamine /0.5 µg plasmid DNA).

### Nitrotyrosine treatment

Nitrotyrosine (3-Nitro-L-tyrosine; Sigma Cat No. 851914) was dissolved in 10 mM HCl to prepare a 10 mM stock. Cells were treated with nitrotyrosine by replacing the medium with nitrotyrosine containing medium or otherwise stated. Equivalent quantities of HCl were used in the medium as control. Medium with or without nitrotyrosine was refreshed every 48 h.

### Cell proliferation

Cells were plated at a density of 2 × 10^5^ cells for HEp-2 and TTLL12 stable clones, 1 × 10^5^ per well for A549, PC3, DU145, PWR-1E, and RWPE1 cells, in 2 ml of medium per well in 6 well multidishes. Cells were allowed to attach and grow for 12 h, and then the medium was replaced with medium containing nitrotyrosine in the treated groups and medium containing an equivalent amount of solvent in the untreated groups, and incubated further as per the experimental requirements. Medium was replaced with fresh medium containing nitrotyrosine or solvent every 48 h. At selected time points, supernatant media in the wells were collected and cells were trypsinized with trypsin-EDTA. Supernatant medium combined with trypsinized cells were counted for viable cells using the tryphan blue exclusion method (44). Cell counts of untreated groups were considered as 100 percent growth. The percent growth of the treated cells over untreated cells was calculated.

### SDS-PAGE and western blots

Cells grown in multiwell dishes were washed twice with cold PBS, lysed in Laemmli buffer (62.5 mM Tris-HCl, pH 6.8, 10% glycerol, 2% SDS) supplemented with Complete® protease inhibitors (Roche), and sonicated before protein quantification (DC BioRad Protein Assay Cat No. 500-0114). Samples with equal protein quantities were supplemented with 5% β- mercaptoethanol, heated (95 °C; 12 minutes), size fractionated on 9% SDS-PAGE gels and transferred to nitrocellulose membranes. All Blue Precision Plus Protein Standard (Bio-Rad Cat. No. 161-0373) was used as the protein size markers. Membranes were blocked for 45 minutes in PBS-T-milk (0.05% Tween, 5% non-fat dried milk); incubated with primary antibodies [diluted in PBS-T milk; 3 h at room temperature (RT)], washed (PBS-T) and incubated with secondary antibodies (1 h; RT; diluted in PBS-T milk). Anti-nitrotyrosine (Upstate Cat No. 06-284) was used at 1:7,000, anti α-tubulin DM1A (Sigma) at 1:1,000; anti-glu tub (AbCys S.A. Cat. No. Abc-0101) at 1:2,000, anti TBP (3G3, IGBMC) at 1: 2,000, anti-rabbit IgG conjugated to HRP (Santa Cruz Biotechnologies Cat. No. sc-2004) and anti-mouse IgG (SantaCruz Biotechnologies Cat. No. sc2005) at 1:10,000. For TBP detection, no milk was used for antibody incubations and no Tween was used during blocking and primary antibody incubations. The TTL antiserum (obtained by injecting peptide 358- VFPPPDVEQPQTQPAAFIKL-377 corresponding to human TTL in New Zealand white rabbits) was used at 1:3,000. The TTLL12 antiserum (obtained by injecting peptide 78- EDEAAREVRKQQPNPGNEL-96 corresponding to human placenta TTLL12 sequence in New Zealand white rabbits) was used at 1:3,000. After removal of unbound secondary antibodies, signals were revealed using SuperSignal™ West Femto Maximum Sensitivity Substrate (Pierce). Signals were quantified using Quantity One software.

### High throughput assay to detect α-tubulin nitrotyrosination in cells

#### Microplate preparation

Exponentially growing TTLL12 and TTL clones were used to seed 96-well polystyrene tissue culture plates (BD MICROTEST ^TM^ 96 Cat. No.353072). The cells were seeded at 10,000 cells in 80 µl of medium per well. TTL clones served as the positive control and TTLL12 clones served as the screening line. Cells were allowed to grow for 12 hours before treatment. Nitrotyrosine was added to each well to get a final concentration of 400 µM. Control wells were treated with the equivalent amount of solvent and DMSO 0.1% final concentration. Background control wells did not receive nitrotyrosine and signal produced by these well were considered as noise and subtracted from the nitrotyrosine signals.

#### Test compound addition

Each well was supplemented with 10 µl MEM containing 100 µM test compounds to get a final test concentration of 10µM in the well, immediately after the addition of nitrotyrosine. Bioinactive controls were obtained with treatment corresponding to DMSO alone. The final concentration of DMSO in the well was 0.1%. After drug dispensing, cells were incubated for 18h at 37°C, 5% CO_2_ in a humidified incubator.

#### Cell immunostaining using TMB

**(C-ELISA-HRP)** 18 h post drug addition, the medium from the wells was aspirated and cells were incubated with 100 µl/well of cytoskeleton isolation buffer (90 mM MES pH 6.7; 1 mM EGTA; 1 mM MgCl_2_; 10% (v/v) glycerol, 0.5% (v/v) Triton X-100) for 3 minutes. Cells were fixed for 10 minutes with 3.7% paraformaldehyde in PBS-0.05% Tween-20. After 3 washes with 200 µl PBS ™0.05% Tween-20, cells were incubated for 1 h at RT with anti-nitrotyrosine antibody diluted 1:1800 or anti glu-tub antibody diluted 1:1,000. Cells were sequentially washed with 200 µl of PBS ™0.05% Tween-20, incubated with goat anti-rabbit secondary antibody diluted in PBS-0.05% Tween-20 for 1 h, washed with 200 µl of PBS-0.05% Tween-20, incubation with 100 µl of TMB for 3 minutes, followed by stopping the reaction with 50 µl of 2N H_2_SO_4_. Plates were read at 450 nm.

#### Cell immunostaining using Europium

**(C-ELISA-Eu)** 18 h after drug addition, the media in the wells were aspirated and cells were incubated with 100 µl/well of cytoskeleton isolation buffer (90 mM MES pH 6.7; 1 mM EGTA; 1 mM MgCl_2_; 10% (v/v) glycerol, 0.5% (v/v) Triton X-100) for 3 minutes 37°C. Cells were fixed for 10 minutes with 3.7% paraformaldehyde in PBS-0.05% Tween-20. After 3 washes with 200µl PBS ™0.05% Tween-20, cells were incubated for 1 h at RT with anti nitrotyrosine antibody diluted 1:1800 in PBS-0.05% Tween-20. Cells were washed with 200 µl of PBS-0.05% Tween-20, and then incubated with Europium labelled secondary antibody diluted 1:1,000 (or otherwise as stated), in PBS-0.05% Tween-20, 1% BSA for 30 minutes. Cells were washed 5 times with 200 µl of PBS-0.05% Tween-20 followed by incubation with 100 µl per well of DELPHIA enhancement solution (Tris-HCl buffered NaCl solution pH 7.8, containing < 0.1% NaN3, bovine serum albumin, bovine gamma globulins, Tween 40, diethylenetriaminepentaacetic acid, and an inert red dye, Perkin Elmer) for 5 min at RT. Plates were read using a time-resolved fluorescence plate reader using 340 nm excitation filter and 615 nm emission filter, with a delay of 400 µs (Victor3, Perkin Elmer).

#### Results analysis

Raw data were exported from the software of the micro plate reader to the Excel file and calculated according to the following procedure. Signals corresponding to nitrotyrosine were obtained by subtracting the noise from cells not treated with nitrotyrosine. Fold increase in nitrotyrosine incorporation by the test substance was calculated by dividing the values of the signals from treated group, by the control group. Test substances showing an increase in signals corresponding to nitrotyrosinated α-tubulins above 1.33-fold at 10 µM concentration was considered as positive hits.

#### Assay performance

To evaluate the quality of the assay, the Z factor was calculated using the following equation according to Zhang et al., 1999 (45):

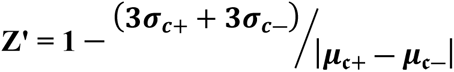

where c+ is the positive control (TTL clones), c- is the screening line (TTLL12 clones), µ is the mean of signal and σ is the SD. In our experimental conditions, the Z factor was above 0.5, indicating that the assay is good for high-throughput screening.

##### Test compound library

A library of 10,000 compounds, called the world biological diversity set, was provided by SPECS (Delft, NL).

## Results

### TTLL12 inhibits nitrotyrosine incorporation into α-tubulin

We previously showed that hTTLL12 affects tyrosination of α-tubulin in cells (10), suggesting that it may also affect nitrotyrosination of α-tubulin (Fig 1). To investigate this possibility, we studied the effects of over- or under-expression of TTLL12 in HEp-2 cells on nitrotyrosination of α-tubulin. The cells were treated with 400 μM of nitrotyrosine for 48 hours, lysates were separated on SDS PAGE and nitrotyrosinated α-tubulin was immunodetected on blots. HEp-2 cell clones that overexpress TTLL12 incorporated less nitrotyrosine into α-tubulin as compared to the control clones (Fig 2A & Fig 2B; two representative clones are shown). Cells in which TTLL12 was downregulated with siRNAs incorporated more nitrotyrosine into α-tubulin as compared to the control siRNAs, for 5 of the 6 siTLL12s tested (Fig 2C & Fig 2D). These results show that TTLL12 inhibits nitrotyrosination of α-tubulin.

**Fig 1.**
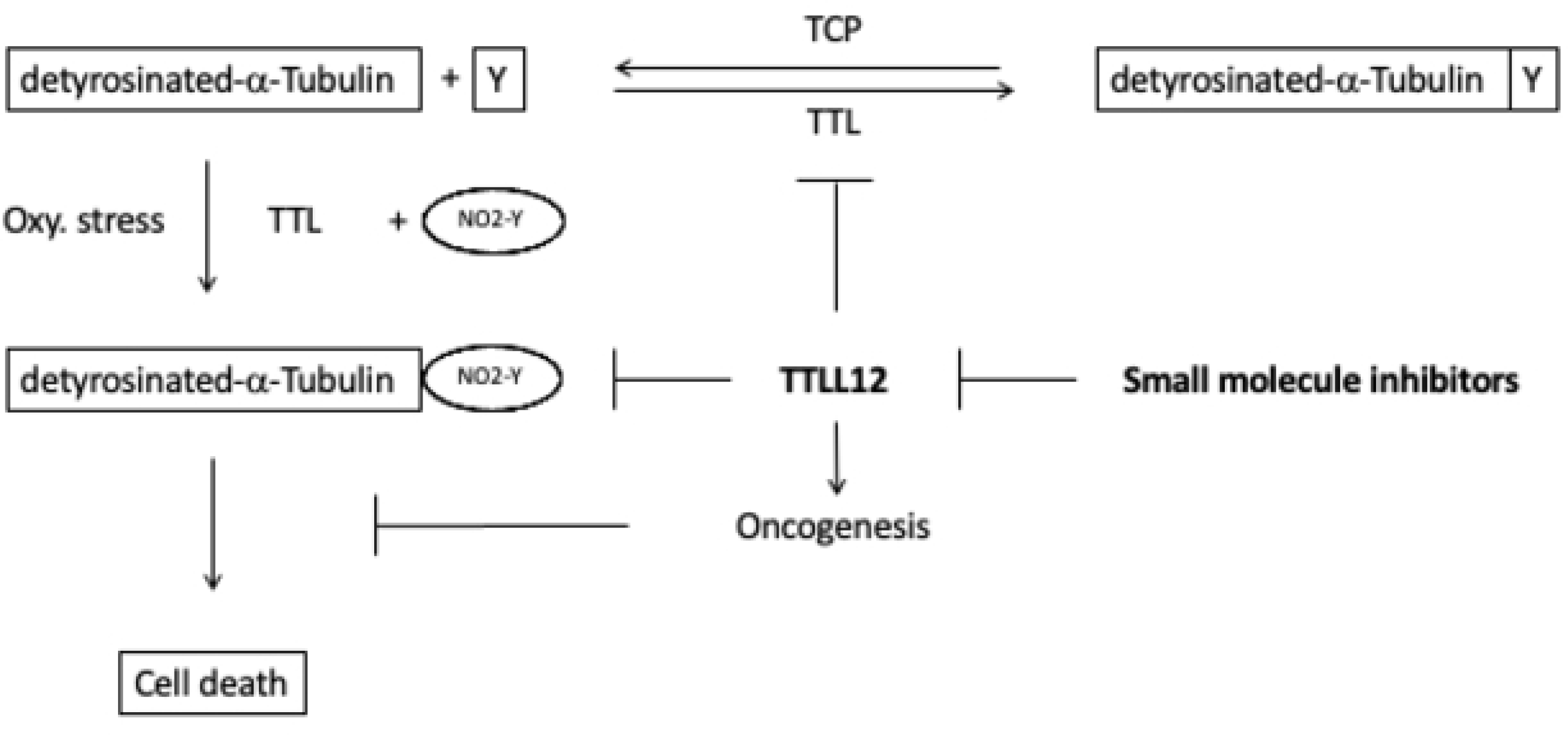
Schematic summary. The diagram illustrates the detyrosination-tyrosination cycle of α-tubulin, in which TCP and TTL catalyse the detyrosination and religation of tyrosine (Y) to α-tubulin. Under oxidative (Oxy.) stress nitrotyrosine (NO2-Y) is ligated to detyrosinated α-tubulin, which has been hypothesised to lead to cell death and suppression of this pathway could lead to oncogenesis. TTTLL12 inhibits ligation of nitrotyrosine to detyrosinated α-tubulin, which is a potential oncogenic activity. Inhibition of this process by small molecule inhibitors could lead to novel therapeutic drugs.

**Fig 2.**
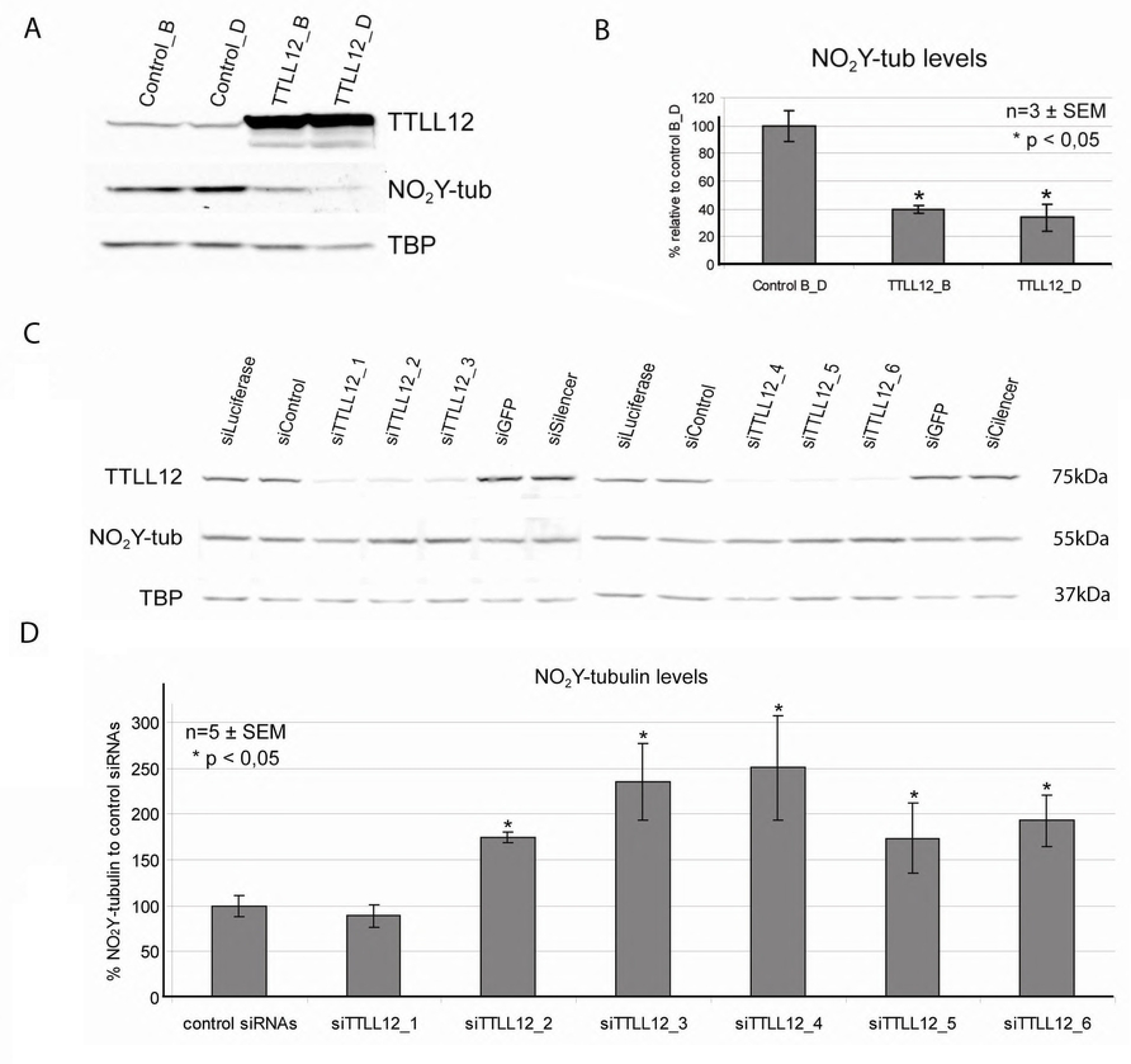
TTLL12 inhibits tubulin nitrotyrosination in HEp-2 cells. (A) Nitrotyrosinated α-tubulin levels in TTLL12 over-expressing clones. The TTLL12 over-expressing clones TTLL12_B and TTLL12_D, and the control clones control_B and control clone_D, were incubated with nitrotyrosine (400 μM) for 48 h. Total lysates were analyzed by immunoblotting with anti-TTLL12, anti-nitrotyrosine and anti-TBP. (B) Densitometric quantification of western blots. Signals corresponding to nitrotyrosine (NO_2_-tub) were normalized to anti-TBP. The average values of nitrotyrosinated tubulin levels of control clones_B and D are presented as 100%. Error bars represent ±SEM of three independent experiments. The symbol * indicates p values < 0.05 calculated with the Student’s t-test. (C) Nitrotyrosinated α-tubulin levels in TTLL12 depleted cells. HEp-2 cells were transfected with 10 nM siRNAs targeting TTLL12 (_1 to _6) or controls (siLuciferase, siControl, siGFP or siSilencer) and treated with nitrotyrosine for 48 h post-transfection. Total cell lysates were analyzed by immunoblotting with anti-TBP, anti-TTLL12 and anti-nitrotyrosine antibodies. Two blots are shown side by side; the control siRNAs are the same on both blots, whereas the siRNAs that target TTLL12 are different. (D) Densitometric quantification of western blots. Signals corresponding to nitrotyrosine were normalized to TBP. Nitrotyrosinated α-tubulin levels of siTTLL12 samples are presented as % of the average of control siRNAs. Error bars represent ± SEM of two independent experiments. The symbol * indicates p < 0.05 calculated with the Student’s t-test, comparing siTTLL12 to control siRNAs.

### Nitrotyrosine inhibits growth of transformed cells

There are contradictory reports of the effects of nitrotyrosine on cell growth [(34, 36)]. Using conditions in which TTLL12 alters nitrotyrosination, we found that nitrotyrosine inhibits the growth of all of the cell lines tested, although there are differences in sensitivity. With 400 μM nitrotyrosine, growth of A549 cells decreased starting from 96 h (Fig 3A), whereas HEp- 2 cells decreased earlier, starting from 72 h, and was lower after 96 h (Fig 3B). The effect of nitrotyrosine on cell proliferation was dose-dependent, since decreasing the amount of nitrotyrosine resulted in smaller effects on cell growth (Fig 3B) as well as lower incorporation into α-tubulin (Fig 3C), showing that there is a correlation between decreased cell growth and nitrotyrosination. To investigate whether transformation may affect sensitivity, we compared two immortalized lines derived from prostate epithelial cells (PWR-1E and RWPE-1) with two lines derived from prostate cancer metastases (DU145 and PC3). The cells were grown in a common medium, to exclude effects of medium composition, such as tyrosine levels. The two metastasis derived lines (DU145 and PC3) were more sensitive to nitrotyrosine, suggesting that transformation and cancer progression could increase sensitivity to nitrotyrosine toxicity (Fig 3D).

**Fig 3.**
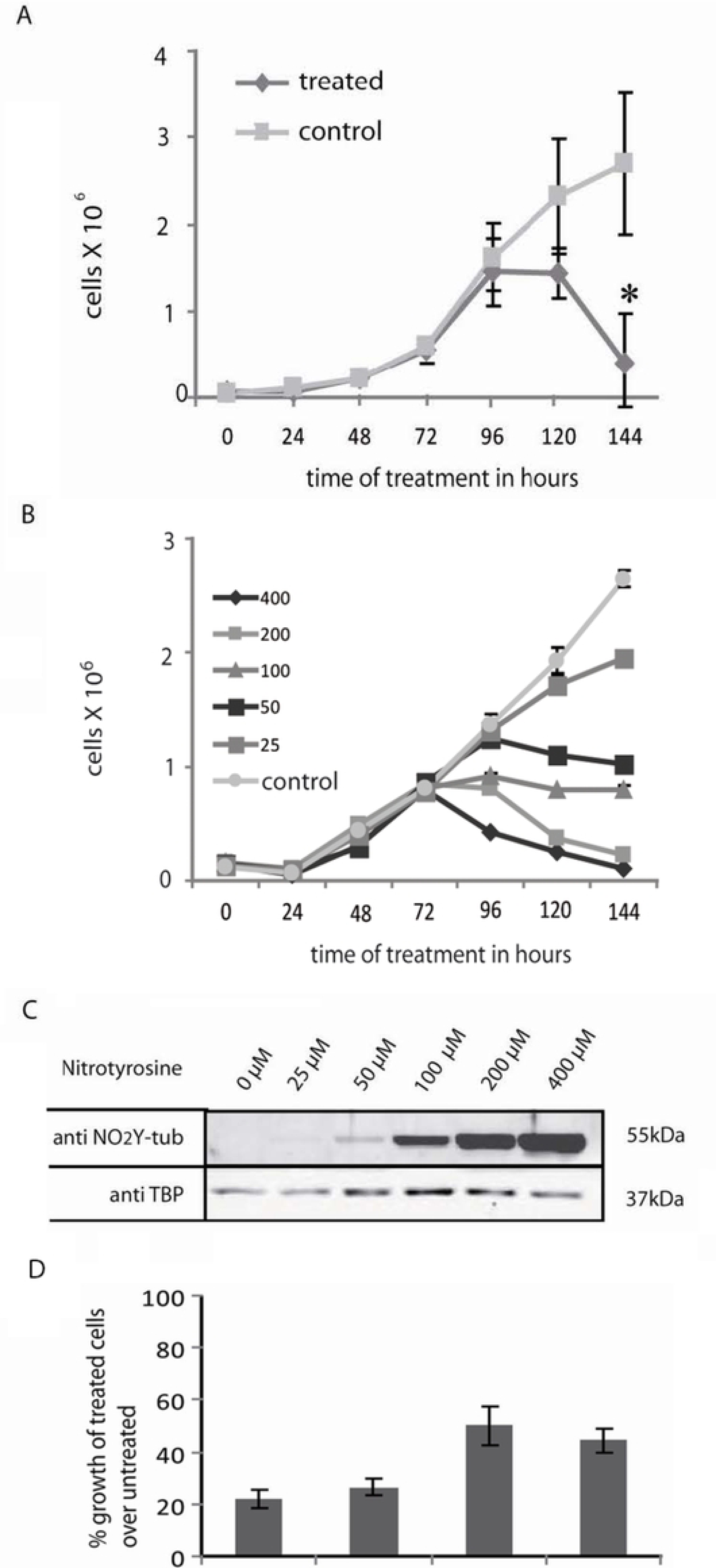
**Nitrotyrosine inhibits cell growth**. (A & B) Cytotoxicity of nitrotyrosine. A549 (A) or HEp-2 (B) cells were cultured in normal medium (control) or in medium supplemented with nitrotyrosine (A, 400 μM; B, 25-400 μM; control cells received the HCl solvent for nitrotyrosine) for 144 h. Media were replaced with corresponding fresh media after 72 h. Viable cells were counted using the tryphan blue exclusion assay and plotted against time. Values represent cells per well and the error bars represent the ± SEM of two independent experiments. p values were calculated with the Student’s t-test, comparing treated with control samples at any particular time point. The symbol * signifies a p value < 0.05. (C) Nitrotyrosinated α-tubulin levels. HEp-2 cells were cultured in medium supplemented with nitrotyrosine (25-400 μM; control cells received HCl used as solvent for nitrotyrosine) for 120 h. Cells were replenished with respective fresh mediums after 72 h. Total cell lysates were analyzed by immunoblotting with anti-nitrotyrosine and anti-TBP antibodies. A representative blot is shown. (D) Cytotoxicity of nitrotyrosine for prostate cell lines. DU145, PC3, PWR1E and RWPE1 cells were cultured for 168 h in a 1:1 combination of media for PC3 and RWPE1 supplemented with either 400 μM of nitrotyrosine or solvent alone. Media were changed every 48 h. Viable cells were counted using the tryphan blue exclusion assay. Values represent % growth of treated cells normalized to untreated controls. The error bars represent the ± SEM of two independent experiments.

### TTLL12 decreases the cell toxicity of nitrotyrosine

Our observation that TTLL12 inhibits nitrotyrosination of α-tubulin suggests that it could decrease the toxicity of nitotyrosine. To test this possibility, we incubated TTLL12 overexpressing and control clones in 50 μM of nitrotyrosine or control medium for 96 h, and then counted viable cells. We found that the TTLL12 over-expressing clones were significantly less sensitive to nitrotyrosine toxicity compared to control clones (Fig 4A). As expected (see above), nitrotyrosination of α-tubulin was found to be lower in TTLL12 over-expressing clones compared to the control clones, whereas the levels of TTL were similar (Fig 4B). In the converse experiment, we found that TTLL12 knockdown decreased the growth of HEp-2 cells in the presence of nitrotyrosine (Fig 5A) and increased nitrotyrosine incorporation into α-tubulin (Fig 5B). We also tested whether DU145, which are relatively sensitive to nitrotyrosine toxicity, could be further sensitized by TTLL12 knockdown. We found that inhibiting TTLL12 expression with siRNA increased the sensitivity of DU145 to nitrotyrosine (Fig 5C) and augmented the incorporation of nitrotyrosine into α-tubulin (Fig 5D). Taken together these results show that TTLL12 expression decreases nitrotyrosine toxicity.

**Fig 4.**
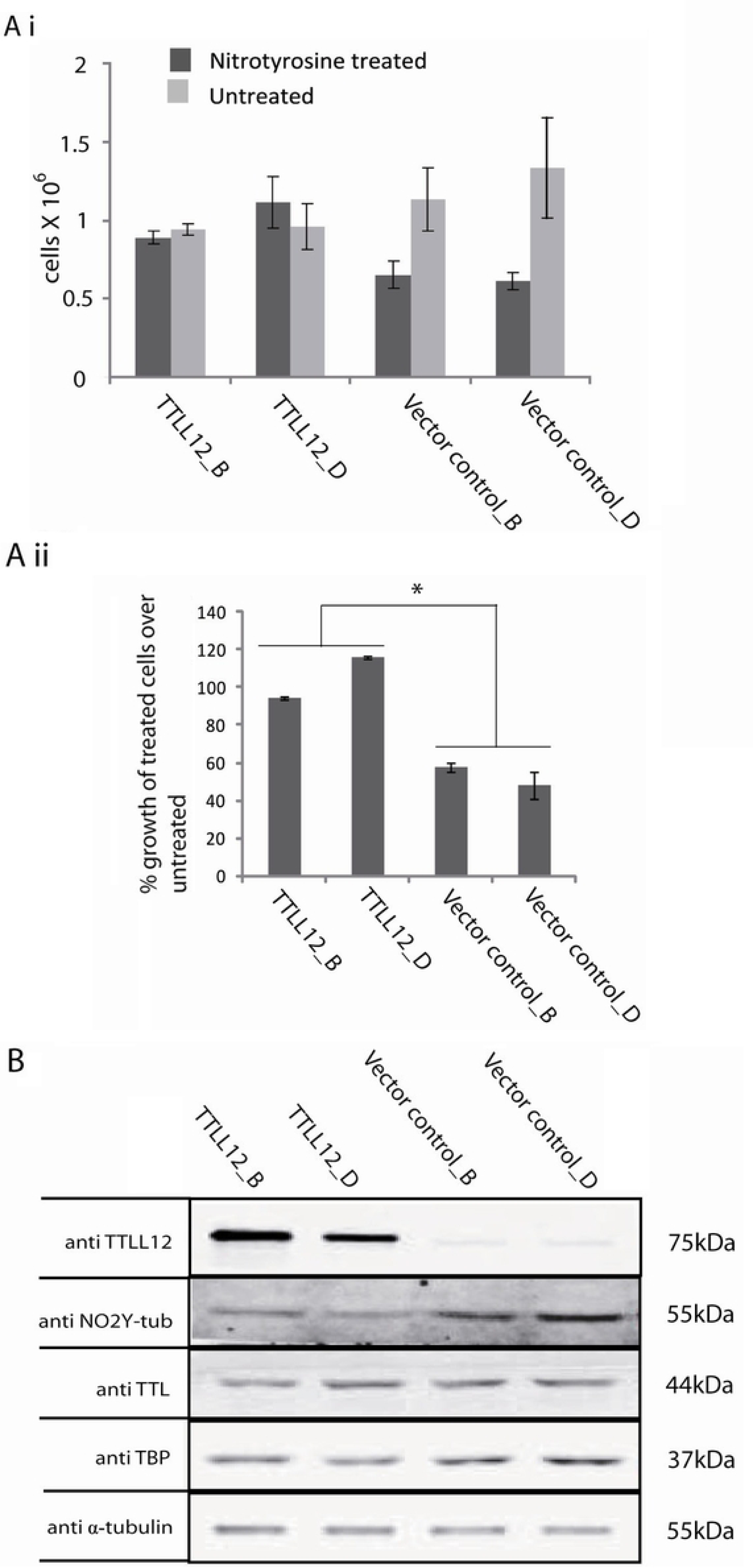
Stable over-expression of TTLL12 in HEp-2 cells increases cell survival in the presence of nitrotyrosine. (A) Cell survival in the presence of nitrotyrosine. HEp-2 cell clones that over-express TTLL12 (TTLL12_B and TTLL12_D) and control clones (Vector control_B and Vector control_D) were cultured for 96 h in medium supplemented with nitrotyrosine (50 μM) or the equivalent amount of solvent. Viable cells were counted using the tryphan blue exclusion assay. Values represent cells per well in the nitrotyrosine treated and untreated groups (i) or % growth of treated cells normalized to untreated controls (ii). The error bars represent the ± SEM of two independent experiments done in duplicates. p values were calculated with the Student’s t-test. The symbol * indicates p values < 0.05. (B) Nitrotyrosinated α-tubulin levels. Total lysates of clones that overexpress TTLL12 and control clones treated with 50 μM nitrotyrosine for 96 h were analyzed by immunoblotting with anti-TTLL12, anti-nitrotyrosine, anti-TTL, anti-TBP, anti-α-tubulin antibodies. A representative immunoblot is shown.

**Fig 5.**
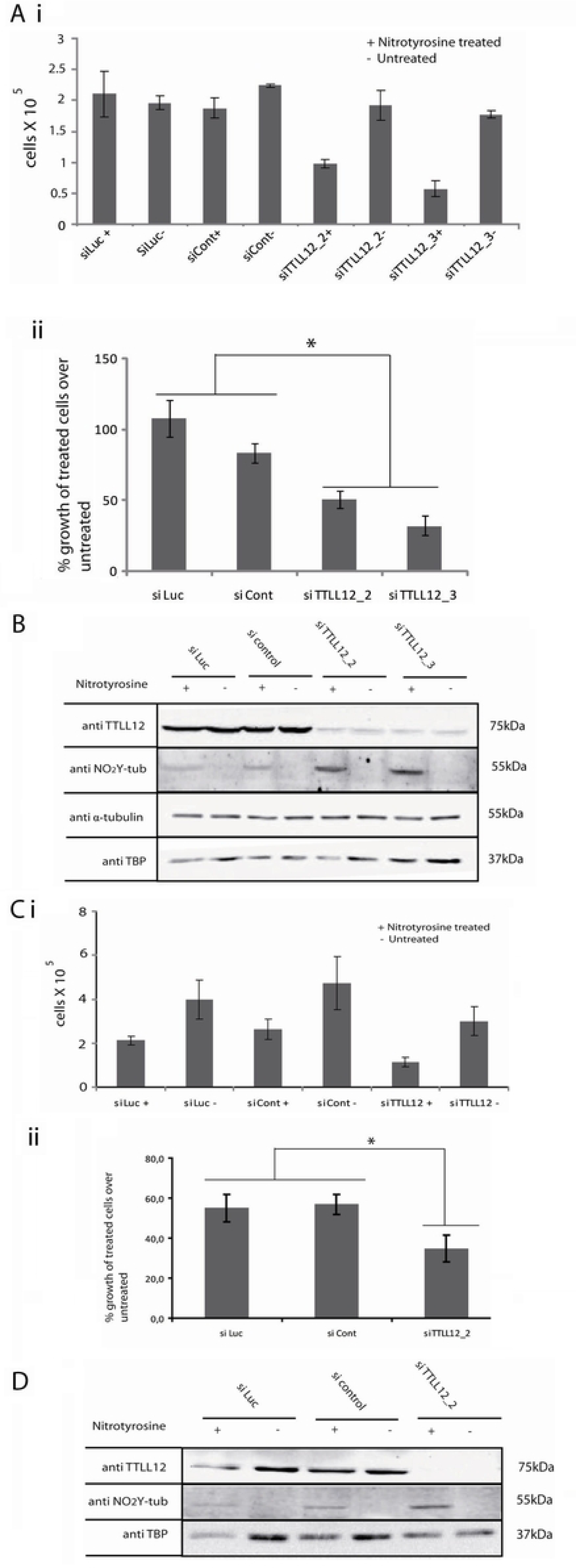
TTLL12 knockdown in HEp-2 and DU145 cells decreases cell survival in the presence of nitrotyrosine. (A) Effect of nitrotyrosine on cell survival. HEp-2 cells were transfected with 10 nM siTTLL12 or control siRNAs and treated for 96 h with 50 μM nitrotyrosine. Controls received medium supplemented with solvent alone. Viable cells were counted using the tryphan blue exclusion assay. (i) Number of cells per well. (ii) Percent growth of treated cells normalized to untreated controls. The error bars represent ± SEM of two independent experiments done in duplicates; p values were calculated with the Student’s t-test. The symbol * indicates p < 0.05. (B) Nitrotyrosinated α-tubulin levels. Using the same culture and treatment conditions as in (A), total cell lysates were prepared and analyzed by immunoblotting with anti-TTLL12, anti-nitrotyrosine, anti-α-tubulin and anti-TBP antibodies. A representative blot is shown. (C) Effect of nitrotyrosine on cell survival. DU145 cells in standard DU145 medium (see Materials and Methods) were transfected with siRNAs targeting TTLL12 or control siRNAs, and treated with 800 μM nitrotyrosine for 96 hours. Control samples received medium supplemented with an equivalent amount of solvent. Viable cells were counted using the tryphan blue exclusion assay. (i) Values represent cells per well in nitrotyrosine treated and untreated samples. (ii) Values represent % growth of nitrotyrosine treated cells normalized to controls. The error bars represent the ± SEM of two independent experiments done in duplicates, p values were calculated with the Student’s t-test (* indicates p < 0.05). (D) Effect of TTLL12 depletion in DU145 on nitrotyrosinated α-tubulin levels. Using the same culture and treatment conditions as (C), total cell lysates were prepared and analyzed by immunoblotting with anti-TTLL12, anti-nitrotyrosine and anti-TBP antibodies. A representative blot is shown.

### Tubulin poisons under non-saturating conditions do not affect nitrotyrosination of α-tubulin

Cytotoxicity induced by microtubule polymerizing and depolymerizing agents could affect nitrotyrosine cytotoxicity by altering incorporation into α-tubulin. We investigated this possibility with the polymerizing agent, paclitaxel, and the depolymerizer, nocodazole, under non-saturating conditions. HEp-2 cells were treated with 100 μM nitrotyrosine, 100 nM paclitaxel, or a combination of the two, for 96 h. Cell growth was progressively inhibited by both molecules and their combination (Fig 6A). Nitrotyrosination of α-tubulin was not affected by paclitaxel at time points at which it was detectable (72 h or later; Fig 6B). Likewise, when HEp-2 cells were treated with nocodazole (1 nM or 10 nM), nitrotyrosine (100 μM) or a combination of the two, cell growth was inhibited. Interestingly, there was a cumulative effect on cell growth, especially after 72 h (Fig 6C). Nitrotyrosination was not affected by 10 nM nocodazole at this time point (72 h), as well as later (Fig 6D). These results show that the cytotoxic effects of nitrotyrosine and tubulin poisons can be cumulative, and suggest that the mechanisms could be different under limiting conditions. Encouragingly, for therapeutic applications, this suggests that manipulating nitrotyrosination could be used in addition to some tubulin poisons, since it would have an additional effect.

**Fig 6.**
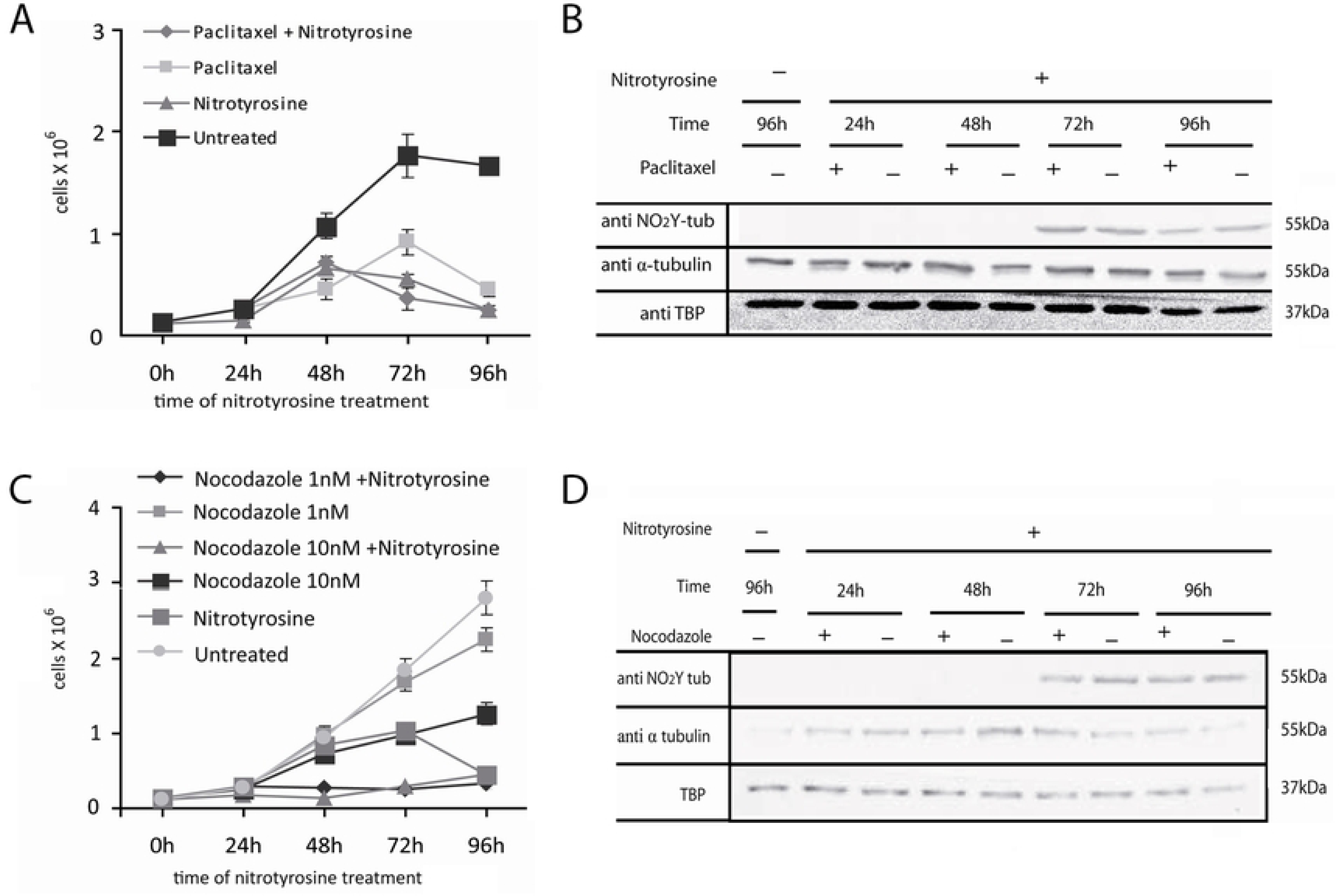
Effect of paclitaxel and nocodazole on cell viability and α-tubulin nitrotyrosination. (A) Growth curves with paclitaxel. HEp-2 cells were treated with nitrotyrosine (100 μM), paclitaxel (100 nM), both or neither (untreated). Viable cells were counted using the tryphan blue exclusion assay and plotted as cells per well against time. The error bars represent the ± SEM of two independent experiments with duplicates in each experiment. (B) Nitrotyrosinated α-tubulin levels with paclitaxel. HEp-2 cells were treated as in A. Total cell lysates were made at the stated times and analyzed by immunoblotting with anti-nitrotyrosine, anti-α-tubulin and anti-TBP antibodies. A representative immunoblot is shown. (C) Growth curves with nocodazole. HEp-2 cells were treated with nitrotyrosine (100 μM), nocodazole (1 or 10 nM), both or neither (untreated, with the equivalent amount of solvents). Viable cells were counted using the tryphan blue exclusion assay and cells per well plotted against time. The error bars represent the ± SEM of two independent experiments with duplicates in each experiment. (D) Nitrotyrosinated α-tubulin levels with nocodazole. HEp-2 cells were treated as in A. Total cell lysates were made at stated time intervals and analyzed by immunoblotting with anti-nitrotyrosine, anti-TBP or anti α-tubulin antibodies. A typical immunoblot is shown.

### Validation of a cell-based screen to measure increases in nitrotyrosination above TTLL12-suppressed levels

The results described above indicate that TTLL12’s suppression of nitrotyrosination is a promising target for the isolation of therapeutic drugs. Consequently, we searched for compounds that could increase nitrotyrosination of α-tubulin in cells that overexpress TTLL12. We used a cell-based ELISA (C-ELISA-HRP), in which cells can be treated with test compounds in the presence of nitrotyrosine and incorporation can be measured directly. Since there are no known inhibitors of TTLL12 and no compound that can increase tubulin nitrotyrosination, we used cell lines that incorporate high or low levels of nitrotyrosine, due to the stable expression of TTL or TTLL12, respectively, to provide a suitable detection window. The assay was adapted from the protocol published by Fonrose et al. (46) and uses a HRP conjugated secondary antibody and a TMB substrate for colorimetric detection. We optimized the assay for reproducibility and linearity, in order to achieve a Z factor (> 0.5) that is suitable for a high throughput screen. The assay effectively detected increases in nitrotyrosinated α-tubulin with increasing time of incubation with nitrotyrosine (S1A Fig). The difference between the high and low nitrotyrosination clones was maintained at all times, with a Z factor > 0.5. The C-ELISA-HRP signal precisely reflected specific bands intensities on immunoblots (S1B Fig). The C-ELISA-HRP efficiently detected the increases in nitrotyrosination that are expected from downregulation of TTLL12 with siRNA (S1C Fig), as well as glu tub (S1D Fig), which could eventually be used in secondary screens. These results validate C-ELISA-HRP for use in screens to select for inhibitors of TTLL12 which increase α-tubulin nitrotyrosination.

We adapted C-ELISA-HRP to a high throughput platform. In particular, the platform uses a time-resolved fluorescence (TRF) detector and a Europium conjugated antibody (C-ELISA- Eu), which provides a higher dynamic range of detection as compared to the colorimetric readout. We optimized the conditions, including antibody dilutions, automated washing steps, time of incubation and buffer conditions (S2 Fig), to maintain Z factors >0.5. Antibodies dilutions of 1:1,000 and 1:2500 were found to be optimal (S2A Fig). Automatic washing effectively replaced manual washing (S2B and C Fig). Times of incubation from 1 to 18 h were suitable for the detection of differences in nitrotyrosination between the high and low nitrotyrosination clones (S2D Fig). The different washing buffers (PBS or Delphia) gave similar values (S2E and F Fig). These results show that the high throughput screening platform (C-ELISA-Eu) gives similar results to the manual assay (C-ELISA-HRP).

### Identification of hits that relieve nitrotyrosination suppression by TTLL12

A library of 10,000 compounds (SPECS^TM^, World Biological Diversity Set) was screened using C-ELISA-Eu on the HTS platform. In the primary screen, nitrotyrosination was increased by 74 compounds (>1.33-fold) and decreased by 16 (<0.38-fold; Fig 7A; only six of the down regulators are show). These hits were rescreened at three different concentrations (10 μM, 1 μM and 0.1 μM). Of the eight selected compounds, six were considered to be of secondary appeal because they were inactive in the secondary screens at 10 μM, or at 1 μM and 0.1 μM (Fig 7B), whereas, interestingly, 2 compounds retained activity at 1 μM (Fig 7C). These two most promising compounds also consistently increased nitrotyrosination of α- tubulin by immunoblotting (Fig 7D), showing that the observed increases are reproducible and specific. These results show that C-ELISA-Eu is a reliable approach for the isolation of specific activators of nitrotyrosination of α-tubulin.

**Fig 7.**
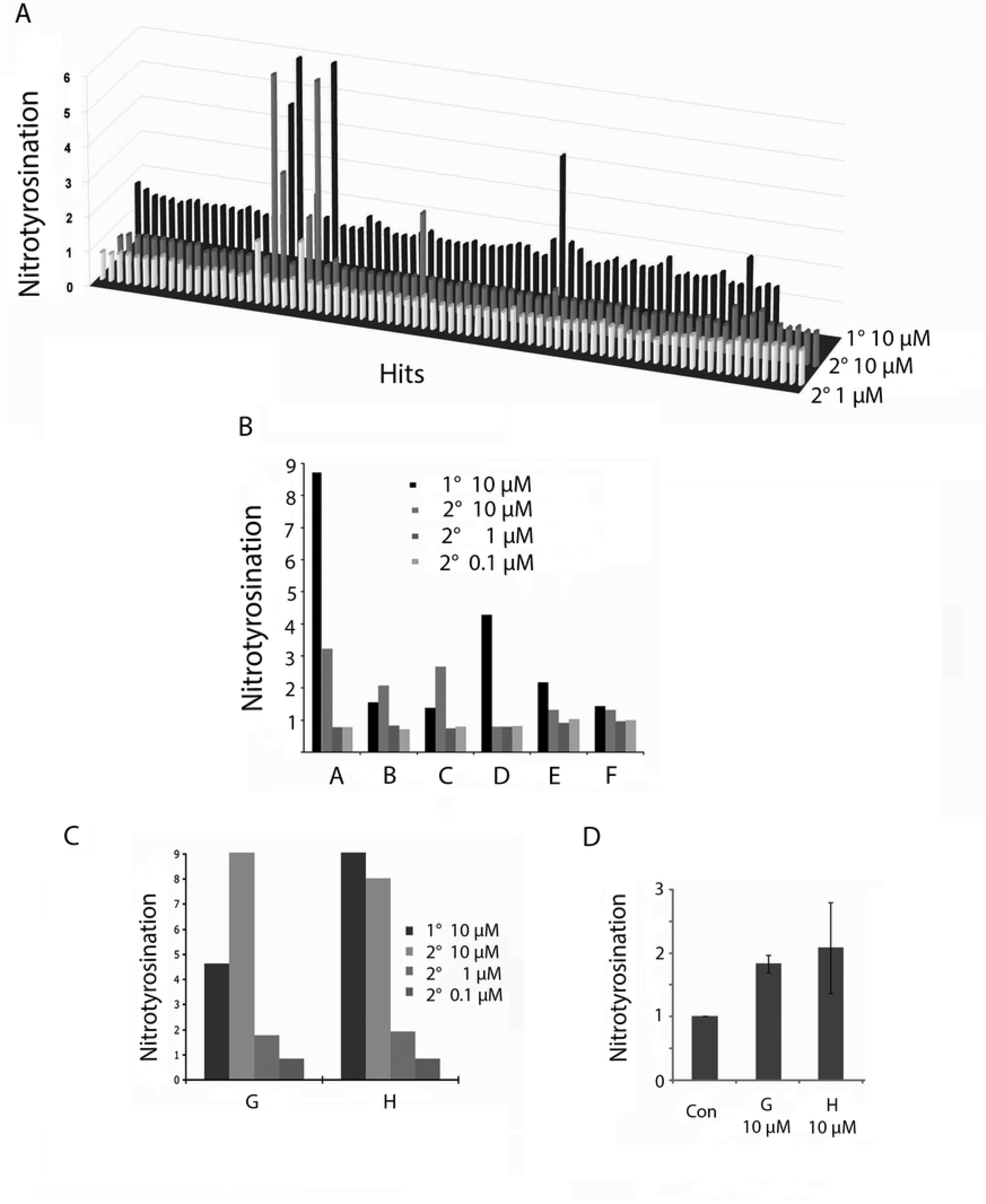
The 10,000-compound screen. (A) 3D graphical representation of fold changes in nitrotyrosine incorporation, focusing on hits that gave fold changes > 1.33 and <0.38 fold. in the primary screen at 10 μM. Secondary screens at 10 and 1 μM, are also shown. (B) Increase in nitrotyrosination induced by six hits, 5 of which retested positive in the 10 μM secondary screen. All were inactive at 1 μM and 0.1 μM. Compound names in library: A = AE- 848/3712516; B = AN-647/1282800; C = AF-399/1503103; D = AP-064/1522801; E = AP-214/1561800; F = AF-399/1503203. (C) Increases in nitrotyrosination induced by the two most promising hits, that were active at both 10 μM and 1 μM in the secondary screen, but not at 0.1 μM. Compound names in library: G = AK-777/12226085; H = AO-289/15480116. (D) Immunoblot verification of the two most promising hits (see C; G = AK-777/12226085; H = AO-289/15480116). TTLL12 clone B cells were treated with 400 μM nitrotyrosine and the indicated concentrations of compounds, and incubated for 18 h. Lysates were analyzed by immunoblotting with anti-nitrotyrosine and anti-α-tubulin antibodies, and the specific signals were quantitated by densitometry. Nitrotyrosinated α-tubulin was normalised to total α-tubulin. The untreated control is considered to be 100%. The error bars represent the ± SEM of two independent experiments with duplicates in each experiment. The y-axes correspond to the fold increase in nitrotyrosination relative to the controls (Con). 1° = primary screen. 2° = secondary screen.

## Discussion

We report a proof of principle that molecules that relieve a potential tumour promoting activity of TTLL12 can be identified, as a step towards developing TTLL12 targeted therapeutic agents (Fig 1). We show that TTLL12 has a potential tumour promoting activity, namely inhibition of the toxicity of nitrotyrosine for transformed cells. Overcoming this activity could be used for therapy, since decreasing TTLL12 levels leads to increased nitrotyrosine toxicity. Manipulating nitrotyrosination of tubulin through TTLL12 appears to be a different microtubule directed therapeutic approach compared to classical tubulin poisons, since they act independently. These findings led us to test whether it was possible to identify molecules that overcome TTLL12 inhibition of nitrotyrosination. We developed a cell-based screen that successfully identified several promising hits from a 10,000 molecule World Biological Diversity Set library. These are original results first reported in a thesis (47).

The ability of TTLL12 to inhibit nitrotyrosine ligation to detyrosinated α-tubulin is consistent with other findings. We previously showed that decreasing TTLL12 levels increased C- terminally detyrosinated α-tubulin levels, similar to reducing TTL levels. However, we were not able to detect the converse changes in C-terminally tyrosinated α-tubulin levels, possibly due to the high endogenous levels of α-tyrosinated tubulin (10). Similar observations have been made with rice TTLL12 (22). Using the tyrosine analogue, nitrotyrosine, we show that there is the expected complementary effect, namely inhibition of nitrotyrosination. Nitrotyrosine is a known substrate of TTL (48) (36), suggesting that nitrotyrosination and tyrosination enter into the same pathways. The mechanism by which TTLL12 inhibits nitrotyrosination remain to be discovered, and may involve protein-protein interactions rather than a direct enzymatic activity. Despite diverse attempts, TTLL12 has not been found to catalyse the various activities predicted from its resemblance to other proteins, (8) (9, 10) (12) suggesting that it is a pseudo-enzyme (10). TTLL12 has been reported to bind α-tubulin (10) (12), indicating that it may physically hinder tyrosine/nitrotyrosine ligation or promote removal.

An intriguing possibility is that TTLL12 could act as a tumour suppressor through suppression of nitrotyrosine ligation to the C-terminus of detyrosinated α-tubulin and the consequent cell toxicity. Idriss (42, 43) proposed that cells have a checkpoint that is triggered by the incorporation of nitrotyrosinated α-tubulin into microtubules, constituting a “last check point” (LCP) that triggers apoptosis. Escape from the LCP leads to cancer. We hypothesized that suppression of nitrotyrosination of α-tubulin by TTLL12 could be one of the mechanisms by which this check point is attenuated. This concords with our and other observations. Nitrotyrosine [reviews: (37) (38)](39-41, 49-52) and TTLL12 (11, 14-19) levels increase with tumour progression, suggesting that they are linked. However, the comparison is indirect, since this nitrotyrosine is mainly found in proteins as a result of chemical modification during oxidative and nitrosative stress, rather than as free nitrotyrosine that is required for ligation to the C-terminus of detyrosinated α-tubulin. The “last check-point” appears to be activatable in transformed cells, since we found that increased nitrotyrosination resulting from decreased expression of TTLL12 was toxic for several transformed cell lines.

Drugs that affect TTLL12 repression of tubulin nitrotyrosination could be redundant with classical tubulin poisons used for therapy since they both impact the tyrosination-detyrosination cycle. Tubulin poisons affect the normal dynamics and stability of microtubules and are thought to thereby affect the extent of tyrosination, since detyrosination occurs on polymerized tubulin, whilst tyrosine is added to free α-β-tubulin dimers by TTL [reviews: (32) (53) (31)]. The effects of tubulin poisons on nitrotyrosine ligation are not well known. We found that the cytotoxicities of tubulin poisons and nitrotyrosine are independent, suggesting that they act through separate mechanism and could be targeted separately for therapy.

Encouraged by these results, we set out to establish if we could develop a robust assay for a high throughput screen to identify activators of TTLL12 suppressed nitrotyrosine incorporation. One of the strengths of our approach is that it is based on nitrotyrosine detection, which is a very good target for the development of assays [see for example: (15) (54) (55) (56)] to follow oxidative stress [review: (57)]. The advantages of our optimized cell-based ELISA are its: compatibility with high throughput screening, sensitivity to detect differences in incorporation with time, capability of detecting increases in tubulin nitrotyrosination induced by inhibition of TTLL12 and usefulness to follow other modifications of tubulin. The assay has a good Z factor, which validates it as a high-throughput screening tool. In cell-based ELISAs test compounds can be tested directly on cells in 96-well plates, whereas sandwich ELISAs require additional steps, cell lysate preparation and protein quantification of the lysates. The current assay could be influenced by indirect effects of the tested molecules on cell division and survival, as well as direct substrate competition with nitrotyrosine.

In order to adapt the assay for a 10,000-molecule screen, a more automated system was developed and optimized. In particular, it uses a time-resolved fluorescence plate reader as the detection system, which has a higher dynamic detection range as compared to the colorimetric readers. The initial assay was developed using HRP conjugated rabbit IgG which provides a colorimetric readout, which was replaced with a Europium conjugated IgG to make the assay amenable to the platform requirements and increasing the dynamic range of detection. Seven compounds were identified that reproducible increase nitrotyrosination. Two of them active when diluted and increased nitrotyrosination in an independent assay (immunoblotting). These results show that C-ELISA-Eu can be used for additional screens and the identified compounds can be further investigated.

To the best of our knowledge, this is the first screen for molecules that modulate nitrotyrosination of tubulin. The molecules from the screen could be used to study the pathways involved in the novel signalling pathway that impinge on modification of the C-terminal tyrosine of α-tubulin, and also nitrosination of tyrosines in proteins. Nitrotyrosination is a novel target for cancer therapy. The molecules from the screen may have their uses in cancer therapy, and also in other pathologies that involve this modification (inflammation, neurodegenerative, and cardiovascular disorders).

## Acknowledgements

We sincerely thank all the members of Procure (FP5), Prima (FP6) and Cancure (FP6) networks, the Amit Deshpande thesis committee, the Wasylyk laboratory members and the IGBMC core facilities for invaluable discussions, suggestions, support and encouragement. The data is this manuscript are original and were first reported in a thesis in 2009 (47). They have since been replicated by a previous member of the laboratory who was aware of these results and worked on other TTLL12 related project in our laboratory (58, 59). BW is an Emeritus Research Director in the CNRS, affiliated to the IGBMC.

## Supporting information

**S1 Fig. Validation of C-ELISA-HRP as a screening assay for increases in nitrotyrosination.**

The optimized cell-based ELISA-HRP detects the difference in α-tubulin nitrotyrosination between TTL (positive control) and TTLL12 clones (screening line). HEp-2 cells were treated with 400 μM nitrotyrosine for 0-24 h and C-ELISA-HRP was performed as described in the Materials and Methods. The error bars represent the ± SEM of three wells in one experiment (* indicates Z factors > 0.5). (B) Western blots confirm the differences in α-tubulin nitrotyrosination and increase with time for both the clones. The TTL and TTLL12 clones were treated with medium supplemented with 400 μM nitrotyrosine for 0-24 h, total lysates were prepared at the indicated times and analyzed by immunoblotting with anti-nitrotyrosine and anti-TBP. (C) C-ELISA-HRP detects increased levels of nitrotyrosinated α-tubulin induced by TTLL12 knockdown. HEp-2 cells were transfected with 10 nM siTTLL12 or si-luciferase and seeded in a 96-well plate. C-ELISA-HRP using nitrotyrosinated tubulin as the readout was performed as described in the Materials and Methods. The Z factor was > 0.5. (D) C-ELISA-HRP detects increased levels of glu-tubulin induced by TTLL12 knockdown. HEp-2 cells were transfected with siTTLL12 or si-luciferase and seeded in 96-well plates. C-ELISA-HRP using glu-tubulin as the readout was performed as described in the Materials and Methods. Z factor = 0.3.

**S2 Fig. Translation of C-ELISA-HRP to the HTS platform.**

Europium conjugated secondary antibodies can be used for the cell-based assay. C-ELISA was performed with Europium conjugated rabbit IgG at different dilutions. The error bars represent the ± SEM of three wells in one experiment (*Z factor > 0.5). (B, C) The washing steps were done either manually (B) or with an automated wash system (C). The error bars represent the ± SEM of three wells in one experiment (* Z factor > 0.5). (D) C-ELISA-Eu detects the increase in α-tubulin nitrotyrosination with time. The error bars represent the ± SEM of three wells in one experiment (*Z factor > 0.5). (E, F) Washing steps with PBS can be replaced with the Delphia^TM^ buffers used for the C_ELISA-Eu. (E) Washing steps before the addition of rabbit IgG were done with PBS, and DELPHIA^TM^ buffer was used thereafter. (F) All the washing steps were performed with DELPHIA^TM^ buffer. The error bars represent the ± SEM of three wells in one experiment. The y-axis represents the time-resolved fluorescence values at emission/excitation wave lengths of 340/615 nm. The symbol * indicates Z factors > 0.5.

